# Lessons and considerations for the creation of universal primers targeting non-conserved, horizontally mobile genes

**DOI:** 10.1101/2020.03.30.017442

**Authors:** Damon C. Brown, Raymond J. Turner

**Affiliations:** Department of Biological Sciences, University of Calgary, Calgary, Alberta, Canada

## Abstract

Effective and accurate primer design is an increasingly important skill as the use of PCR-based diagnostics in clinical and environmental settings is on the rise. While universal primer sets have been successfully designed for highly conserved core genes such as 16S rRNA and characteristic genes such as *dsrAB* and *dnaJ*, primer sets for mobile, accessory genes such as multidrug resistance efflux pumps (MDREP) have not been explored. Here, we describe an approach to create universal primer sets for select MDREP genes chosen from five superfamilies (SMR, MFS, MATE, ABC and RND) identified in a model community of six members *(Acetobacterium woodii, Bacillus subtilis, Desulfovibrio vulgaris, Geoalkalibacter subterraneus, Pseudomonas putida* and *Thauera aromatica).* Using sequence alignments and *in silico* PCR analyses, a new approach for creating universal primers sets targeting mobile, non-conserved genes has been developed and compared to more traditional approaches used for highly conserved genes. A discussion of the potential shortfalls of the primer sets designed this way are described. The approach described here can be adapted to any unique gene set and aid in creating a wider, more robust library of primer sets to detect less conserved genes and improve the field of PCR-based screening research.

**Importance:** Increasing use of molecular detection methods, specifically PCR and qPCR, requires utmost confidence in the results while minimizing false positives and negatives due to poor primer designs. Frequently, these detection methods are focused on conserved, core genes which limits their applications. These screening methods are being used in various industries for specific genetic targets or key organisms such as viral or infectious strains, or characteristic genes indicating the presence of key metabolic processes. The significance of this work is to improve primer design approaches to broaden the scope of detectable genes. The use of the techniques explored here will improve detection of non-conserved genes through unique primer design approaches. Additionally, the approaches here highlight additional, important information which can be gleaned during the *in silico* phase of primer design which will improve our gene annotations based on sequence homologies.

## Introduction

PCR-based diagnostic approaches are being widely used for rapid screening of microbes and pathogens in environmental and clinical settings (1–4). Correct primer design is a critical factor in assessing the accuracy of the diagnostic approach to avoid false positives and false negatives. Many clinical studies focus on a single infectious strain or species (5–7) and thus the primers can be designed to focus on specific, characteristic genes present in the pathogenic strains and not in the benign strains, simplifying the primer design. However, this restricts the scope of use for the primers.

Some of the most successful attempts at designing “universal” primers are the 16S rRNA sets, of which there are many subsets, each targeting different variable regions of the 16S rRNA gene and are reviewed elsewhere (8–10). These universal primer sets are mixed batches of primers with differing degrees of base variability at each position within the primer, where each possible combination is present and intended to target specific sequences, each representing a different species or groups of species. This approach attempts to cover the depth of diversity within the target location and provide unbiased detection of each species present before the first PCR cycle. Over the subsequent years, it has been shown that different primer sets have varying detection levels of the species, both within different bacterial clades and between *Bacteria* and *Archaea* (8, 11, 12). Thus, the idea of ‘universal’ is difficult even for an expectedly highly conserved core gene.

Here, we describe an approach to design universal primer sets targeting multidrug resistance efflux pumps (MDREPs). MDREPs are non-conserved genes and cover a vast range of different proteins ranging from specific metabolite efflux transporters to those targeting specific groups of antiseptics and/or antibiotics (13–15). They are a group of integral membrane transport protein systems subdivided into six superfamilies: small multidrug resistance (SMR), major facilitator superfamily (MFS), multidrug and toxic (compound) extrusion (MATE), ATP-binding cassette (ABC), resistance-nodulation-cell division (RND) and proteobacterial antimicrobial compound efflux (PACE). A review of the superfamilies is available elsewhere (13). Due to the narrow range of annotated PACE genes (namely *aceI)*, and the absence of any annotated PACE genes present in the genomes of the model community members, PACE genes are omitted from this study.

As their name suggests, MDREPs were originally classified based on their ability to confer resistance to antibiotics, although a single MDREP can have a diverse substrate range, sharing little structural, size or ionic properties (16–18). To date, it is unclear how the specificity of the proteins is determined, however recent research suggests it may be from different entrance channels allowing transport of chemicals with similar physicochemical properties or through the use of weaker hydrophobic interactions between substrate and efflux pump compared to specific hydrogen bonds used by more specific transporters (19, 20).

MDREPs are often found on mobile genetic elements including plasmids, transposons, integrons, integrative conjugative elements and genomic islands (15, 21–24). Thus, they are of particular and increasing importance due to their horizontal mobility and their contribution to the growing problem of global antibiotic resistance (24–26). The phenomenon of antibiotic resistance is well studied in medical environments, but has only recently begun to be investigated in other environments such as water treatment plants and activated sludges (27–29). Several research groups have begun using metagenomics to track and monitor the migration and abundance of different MDREP genes in waste water following various treatment methods to mixed results (30, 31). Our interest is to follow specific MDREPs in the context of biocide resistance in a model community of six members designed to resemble a microbiologically influenced corrosion environment.

Many detailed reviews of the different efflux pump superfamilies have been published (32–37) which have been used for the selection of targets in this study and are shown in Table 1. Here, gene targets have been specifically selected for their published substrate compounds, focusing on antiseptics/biocides.

**Table 1.** Targeted multidrug resistance efflux pump genes targeted in this study and their details

| Superfamily | gene | substrates | References |
| --- | --- | --- | --- |
| MFS | *emrB* | Carbonyl cyanide m-chlorophenylhydrazone (CCCP), tetrachlorosalicyl anilide, nalidixic acid, thioactomycin | (38, 39) |
|  | *qacA* | Ethidium bromide, cetrime, benzalkonium chloride, chlorhexidine | (40) |
| MATE | *norM* | Norfloxacin, doxorubicin, acriflavine | (41) |
|  | *mepA* | Tigecycline, pentamidine, ethidium bromide, dequalinium, tetraphenylphosphonium, 1-(4-trimethylammoniumphenyl)-6-phenyl-1,3,5-hexatriene p-toluenesulfonate, chlorhexidine, norfloxacin and ciprofloxacin | (42, 43) |
| RND | *acrB* | Taurocholate, glycocholate, tetracycline, chloramphenicol, fluoroquinolone, novobiocin, erythromycin, fusidic acid, rifampin, ethidium bromide, acriflavine, crystal violet, sodium dodecyl sulfate, deoxycholate, nafcillin, cloxacillin, | (44–46) |
|  | *mexB* | Quinolones, macrolides, tetracyclines, lincomycin, chloramphenicol, novobiocin, oxacillin, ethidium bromide, tetraphenylphosphonium, sodium dodecyl sulfate | (47, 48) |
| SMR | *ebrAB* | Ethidium bromide, tetraphenylphosphonium chloride, acriflavine, pyronine Y and safranin O | (49, 50) |
|  | *emrE* | Quaternary cation/ammonium compounds (e.g. Betaine, choline, methyl viologen, tetraphenylphosphonium, benzalkonium, cetyltrimethylammonium bromide, cetylpyridinium chloride), ethidium bromide, acriflavine, crystal violet, pyronine Y, safranin O | (51, 52) |
|  | *qacE* | Ethidium bromide, proflavine, rhodamine, cetyltrimethylammonium bromide, benzalkonium chloride, tetraphenylarsonium chloride, diamidinodiphenylamine dichloride, propamidine isethionate, pentamidine isethionate | (40, 53) |
|  | *sugE* | Tributyltin, ethidium bromide, chloramphenicol, tetracycline, cetylpyridinium, cetyldimethylethyl ammonium, cetrime | (54, 55) |
| ABC | *lmrA* | Vinblastine, vincristine, daunomycin, doxorubicin, colchicine, ethidium bromide, valinomycin, nigericin, Hoechst 33342, diphenylhexatriene, rhodamine 6G | (56) |

We explore the possibility to design universal primers for non-conserved, mobile gene targets in the fashion of the multiple universal 16S rRNA primer sets (2, 8). We will discuss difficulties and challenges encountered and describe a novel approach which can be applied to any desired gene target for improved detection. Our group’s focus is on MDREPs that would provide resistance to biocides used in microbiologically influenced corrosion and biofouling control.

## Methods

### *In silico* gene identification

Six bacteria were chosen to represent a highly simplified mixed species community as may be identified in a microbiologically influenced corrosion environment. The chosen strains all had fully sequenced genomes available on the Joint Genome Institute website (https://img.jgi.doe.gov/cgi-bin/m/main.cgi). The IMG Genome ID numbers for the representative genomes are as follows: *Acetobacterium woodii* (2512047040), *Bacillus subtilis* (2511231064), *Desulfovibrio vulgaris* (637000096), *Geoalkalibacter subterraneus* (2593339207), *Pseudomonas putida* (641522645), and *Thauera aromatica* (2791355019).

The full genome sequences were imported from JGI to a web-hosted sequence alignment tool (https://benchling.com/) for gene sequence alignments. Multidrug resistance efflux pump genes were identified through searching directly for their gene names and abbreviations or key words including (but not exclusively): multidrug, efflux, transporter, outer membrane, inner membrane, and resistance. A recognized challenge was that not all genomes were equally annotated. For example, the genome of *P. putida* has a more complete annotation compared to the more environmentally relevant species such as *D. vulgaris.* Once identified, all copies of each gene were clustered and a nucleic acid multiple sequence alignment (MSA) was performed with no template sequence and using the MAFFT alignment algorithm with default conditions (maximum iterations: 0, tree rebuilding number: 2, gap open penalty: 1.53, gap extension penalty: 0.0 and no adjust direction). MSAs were used to identify regions of nucleotide homology across all annotated genes of the same name. These regions were preferentially used for primer design as described below. Gene details were taken from the NCBI GenBank files for each species once the locus tags were identified on Benchling. In cases where no product name was provided for a specific locus tag, the protein ID was investigated and a gene identity was assigned from the region name.

### Primer design approaches

#### Primer design approach A (Traditional)

Using the MSA, homologous regions of the nucleotide sequences were identified and used as targets for primer binding. Primers were all designed to be 18-23 base pairs in length and create PCR amplicons of 180-240 base pairs in length to facilitate quantitative PCR (qPCR) analysis as a downstream application. GC content was targeted to be 50% but exceptions were made which allowed the GC content to reach a maximum of 80% (Table 2) in order to accommodate the target regions. Although efforts were made to maintain a melting temperature of ±5 °C between the upstream (forward) and downstream (reverse) primers for all primer pairs of a specific gene, priority was given to optimizing the melting temperatures of a specific pairing (albeit this rule had to be stretched on occasion as well). The position along the MSA which matched all these conditions were used to create primers, and when required primers were designed with degenerate bases to ensure they targeted the desired locations of the MSA.

**Table 2.**
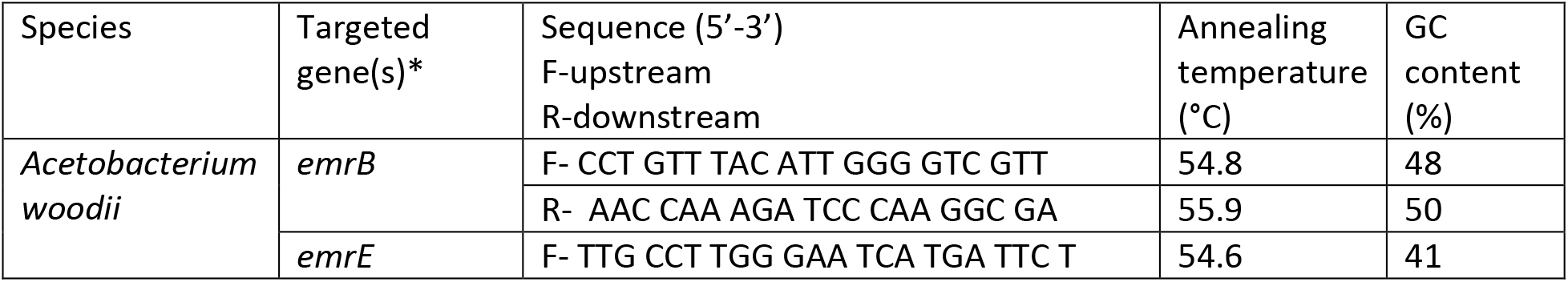
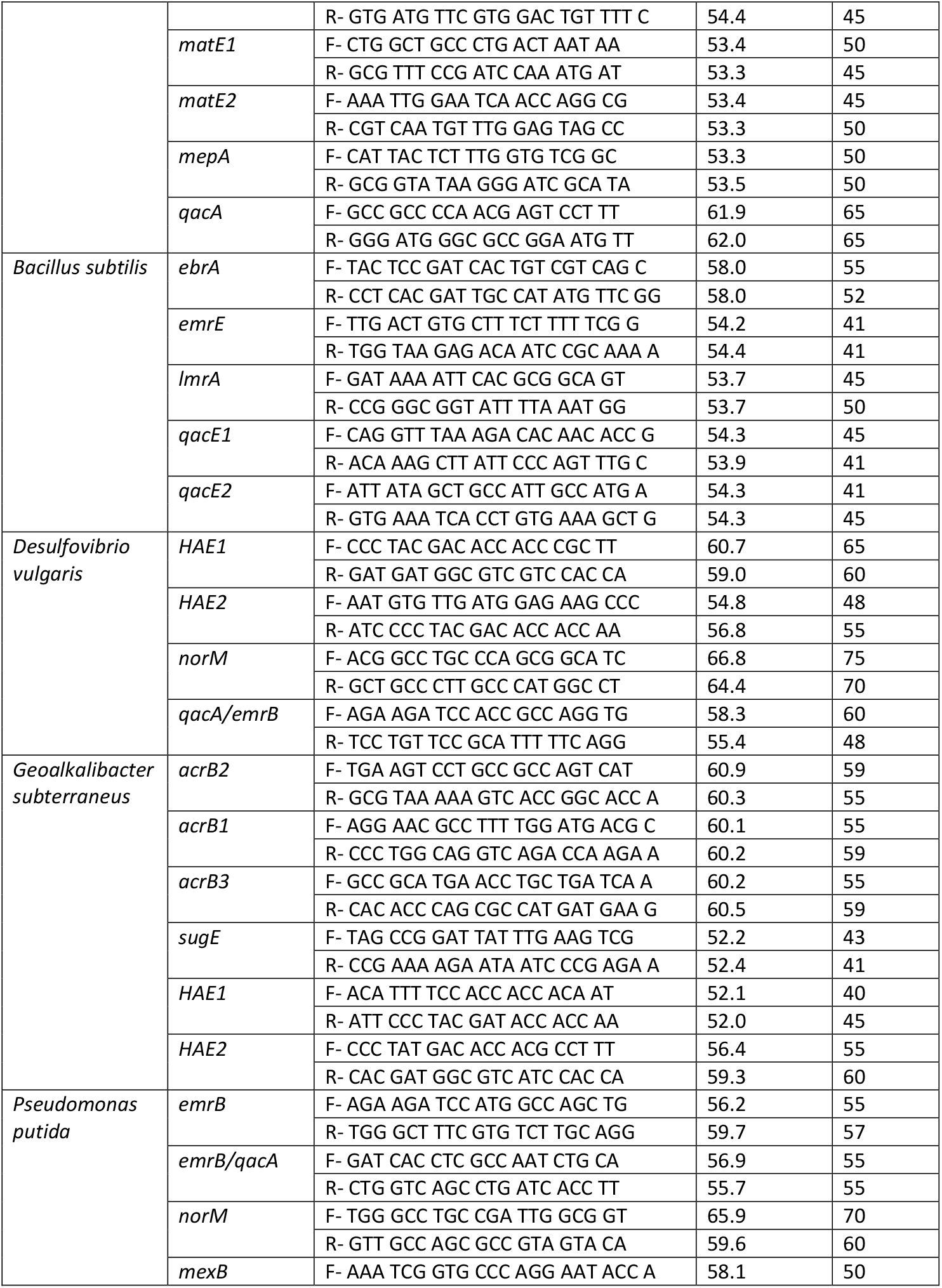
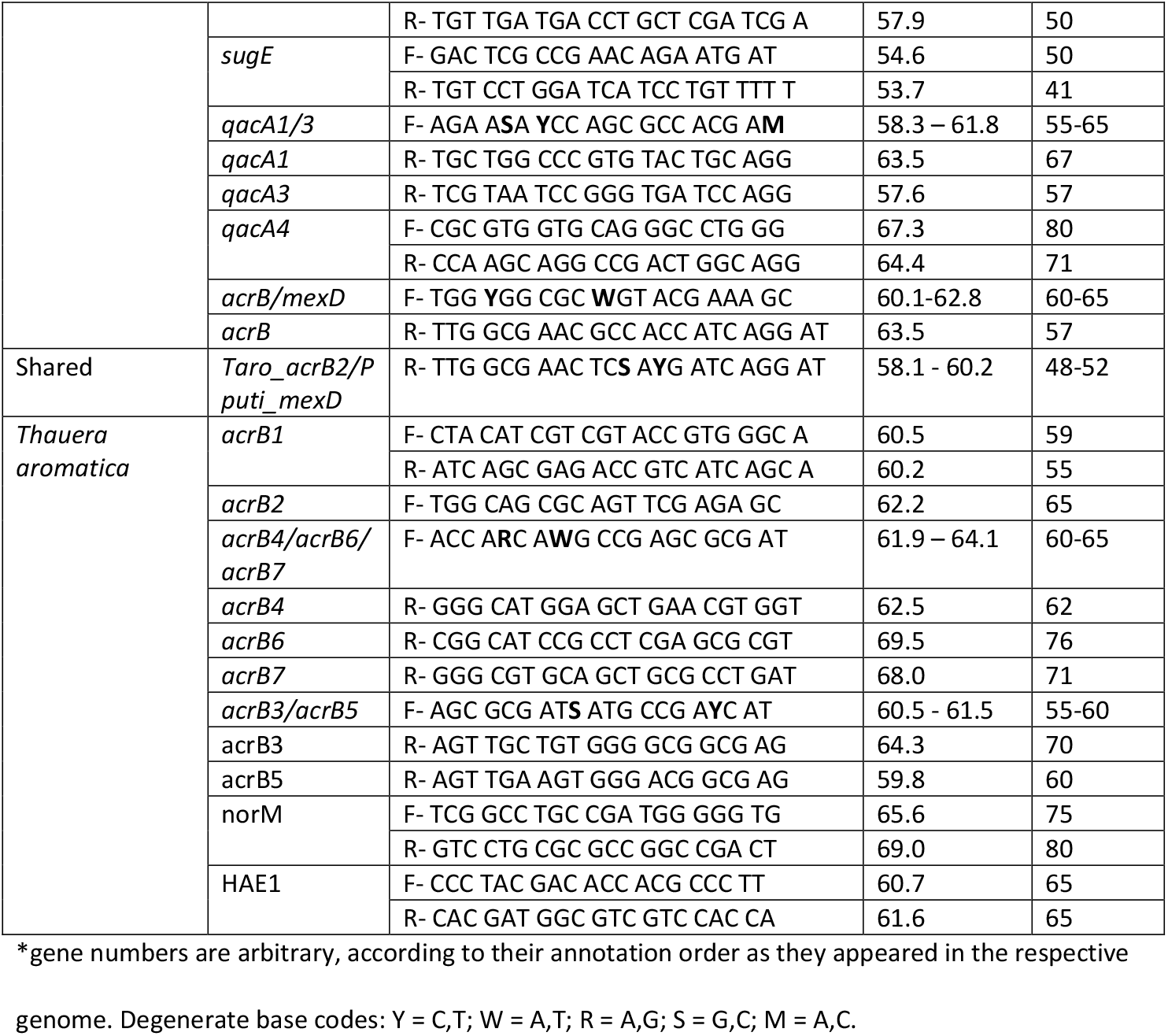
PCR primer details and amplification protocols

#### Primer design approach B (Novel)

As a result of the limitations of the primer design approach A, a unique approach was developed to more efficiently target genes annotated the same but with a lower sequence homology. Primers were designed to be 18-23 nucleotides in length, produce an amplicon of 180-240 base pairs and have the same melting temperatures ± 5 °C between the upstream and downstream primer pairs. To avoid the use of degenerate bases, the primer positioning against the MSA was more flexible, allowing the bind locations for the primer pairs to drift while maintaining identical amplicon size. Using this approach, the amplicon size was a higher priority than annealing location to allow for identical amplicon size to prevent issues with downstream analysis across different primer pairs targeting same genes in different genomes.

### Primer binding testing

Intended and unintended primer binding locations were identified using the following *in silico* conditions on the Benchling platform. Primer binding parameters were a minimum of 18 matched bases, a maximum of three mismatches total with no consecutive mismatches and annealing temperatures between 30-100 °C. Due to the binding algorithm of Benchling’s primer binding, primers with mismatching ends had to be manually removed from the pool (no mismatches were allowed on the primer ends). A full list of the binding locations for all primers is available in Supplementary Table 1.

## Results and Discussion

The goal of this study was to go through a primer design workflow, then test the efficacy of the designed primer sets *in silico* to evaluate the success or failure of the design. This approach is to evaluate the primers before using the primers experimentally, consuming resources and time in optimisation. The output of our *in silico* primer annealing experiment is represented over five figures which illustrate all the binding locations for each of the primers within the model community genomes separated into MDREP super families (Figures 1–5). The results illustrate the direct hits towards the intended MDREP gene but also the unintended binding locations where primers would anneal under different annealing temperatures and with different sequence homologies. In all figures, the type of line used indicates the percent of sequence homology (solid line 100%, dashed line 90-99.9%, dotted line 80-89.9%) and the colour coding is used to indicate the melting temperature range divided into 5 °C increments (light green ≤49.9 °C, orange 50-54.9 °C, blue 55.0-59.9 °C, purple 60.0-64.9 °C, red 65.0-69.9 °C and dark red ≥70.0 °C). For simplicity the term “hypothetical protein” refers to all genes annotated as: conserved protein of unknown function, hypothetical protein, conserved hypothetical protein, conserved protein of unknown function, and conserved exported protein of unknown function.

**Figure 1.**
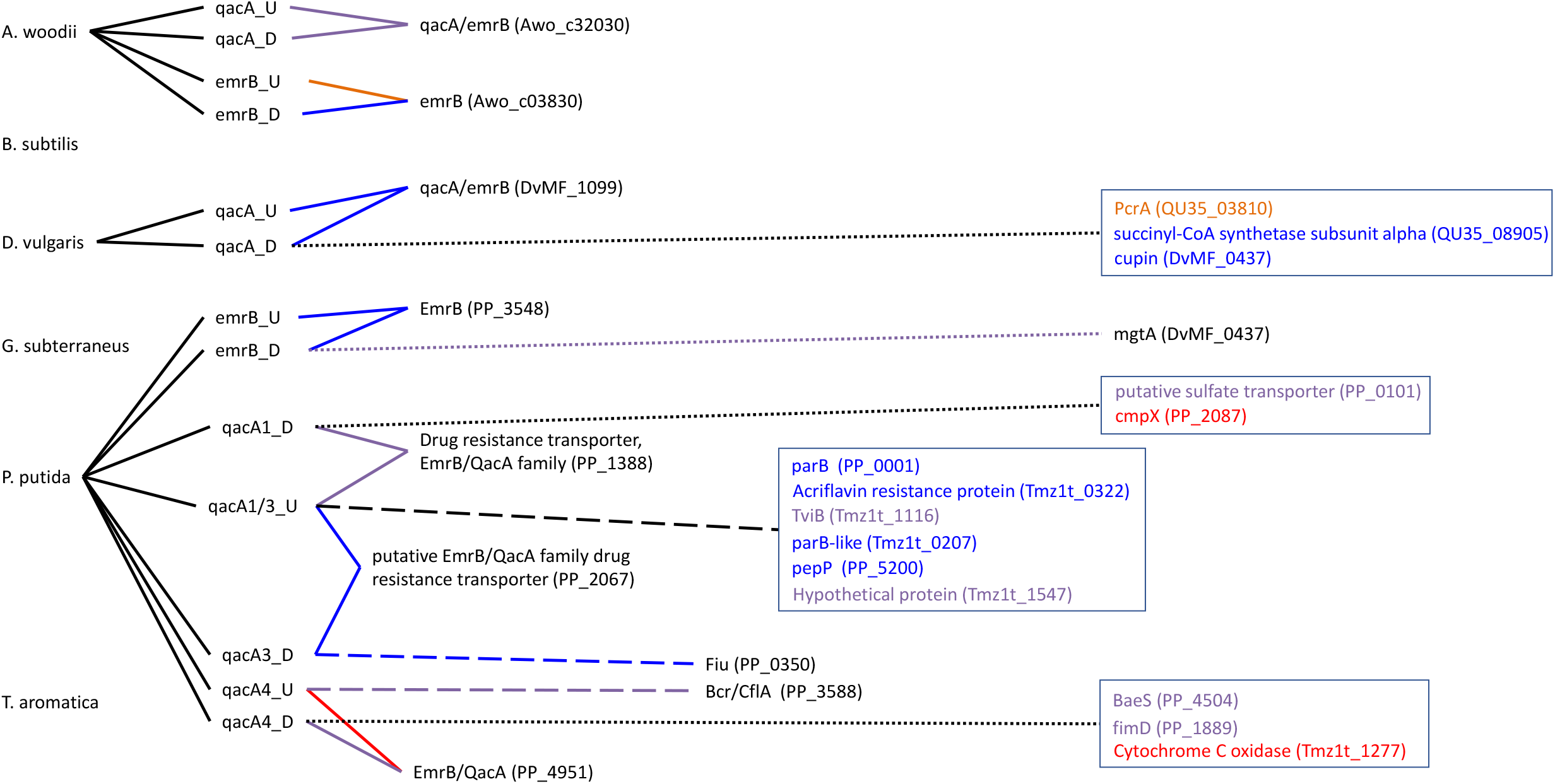
Visual representation of the primers designed in this study targeting the Major Facilitator Superfamily (MFS) genes and all targets they bind to. The type of line connecting the primer and the target indicate the homology of binding (solid line 100%, dashed line 90-99.9%, dotted line 80-89.9%). The colour coding is used to indicate the melting temperature range divided into 5 °C increments (light green ≤49.9 °C, orange 50-54.9 °C, blue 55.0-59.9 °C, purple 60.0-64.9 °C, red 65.0-69.9 °C and dark red ≥70.0 °C). A black line connecting to the gene targets grouped into a box indicates the homology of all genes in the box and the text colour represents the melting temperature.

**Figure 2.**
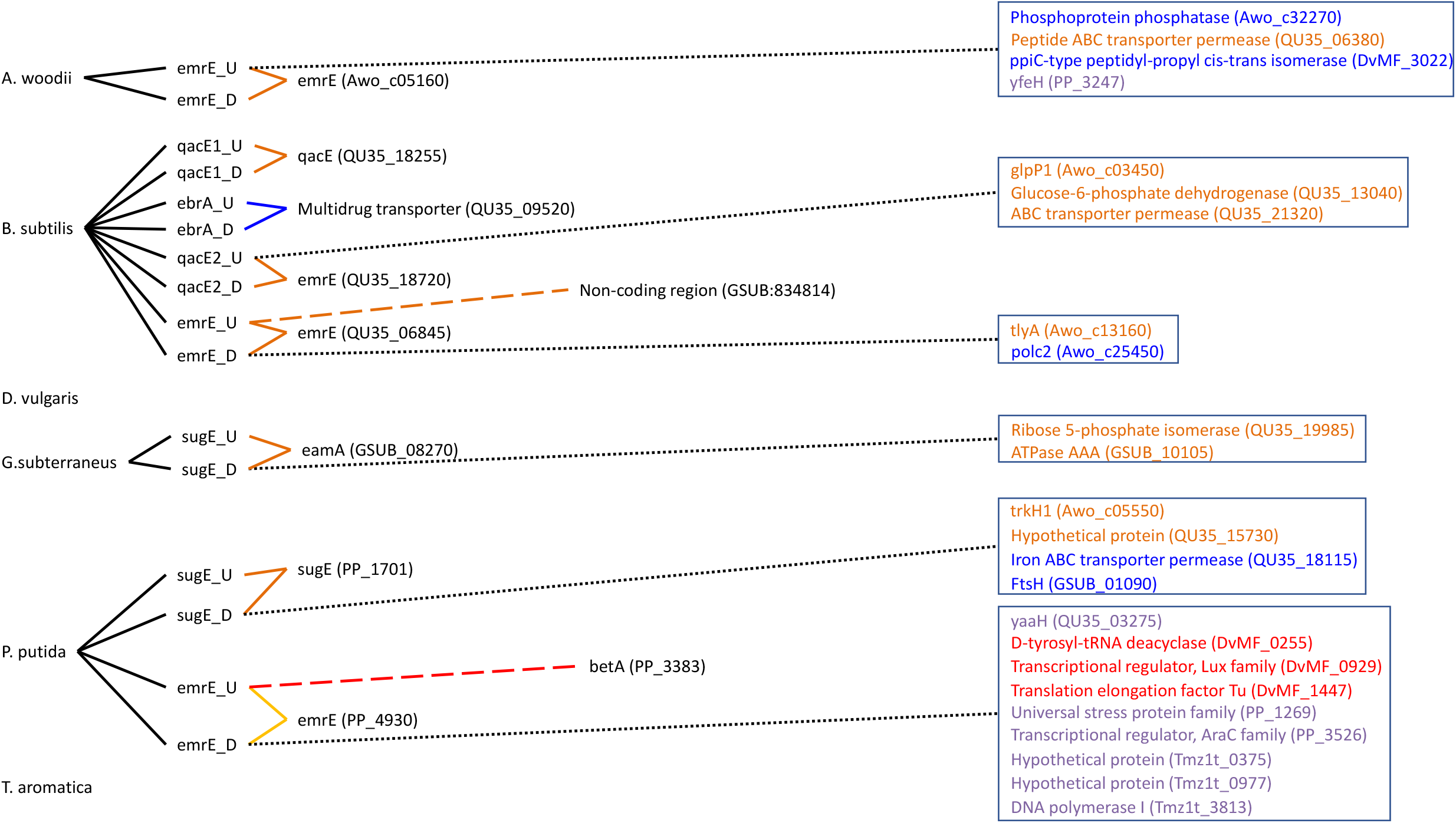
Visual representation of the primers designed in this study targeting the Small Multidrug Resistance (SMR) genes and all targets they bind to. The type of line connecting the primer and the target indicate the homology of binding (solid line 100%, dashed line 90-99.9%, dotted line 80-89.9%). The colour coding is used to indicate the melting temperature range divided into 5 °C increments (light green ≤49.9 °C, orange 50-54.9 °C, blue 55.0-59.9 °C, purple 60.0-64.9 °C, red 65.0-69.9 °C and dark red ≥70.0 °C). A black line connecting to the gene targets grouped into a box indicates the homology of all genes in the box and the text colour represents the melting temperature.

**Figure 3.**
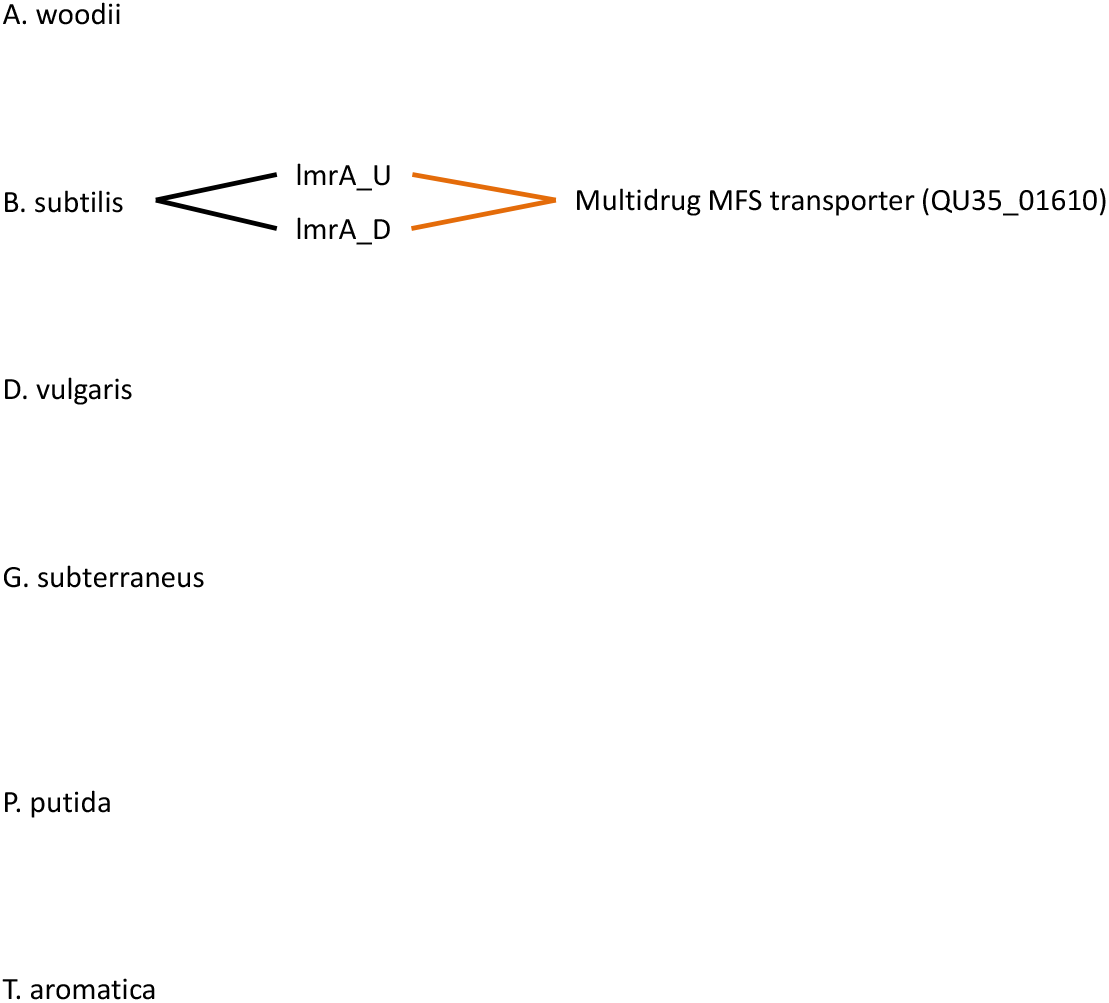
Visual representation of the primers designed in this study targeting the ATP-binding cassette (ABC) genes and all targets they bind to. The type of line connecting the primer and the target indicate the homology of binding (solid line 100%, dashed line 90-99.9%, dotted line 80-89.9%). The colour coding is used to indicate the melting temperature range divided into 5 °C increments (light green ≤49.9 °C, orange 50-54.9 °C, blue 55.0-59.9 °C, purple 60.0-64.9 °C, red 65.0-69.9 °C and dark red ≥70.0 °C). A black line connecting to the gene targets grouped into a box indicates the homology of all genes in the box and the text colour represents the melting temperature.

**Figure 4.**
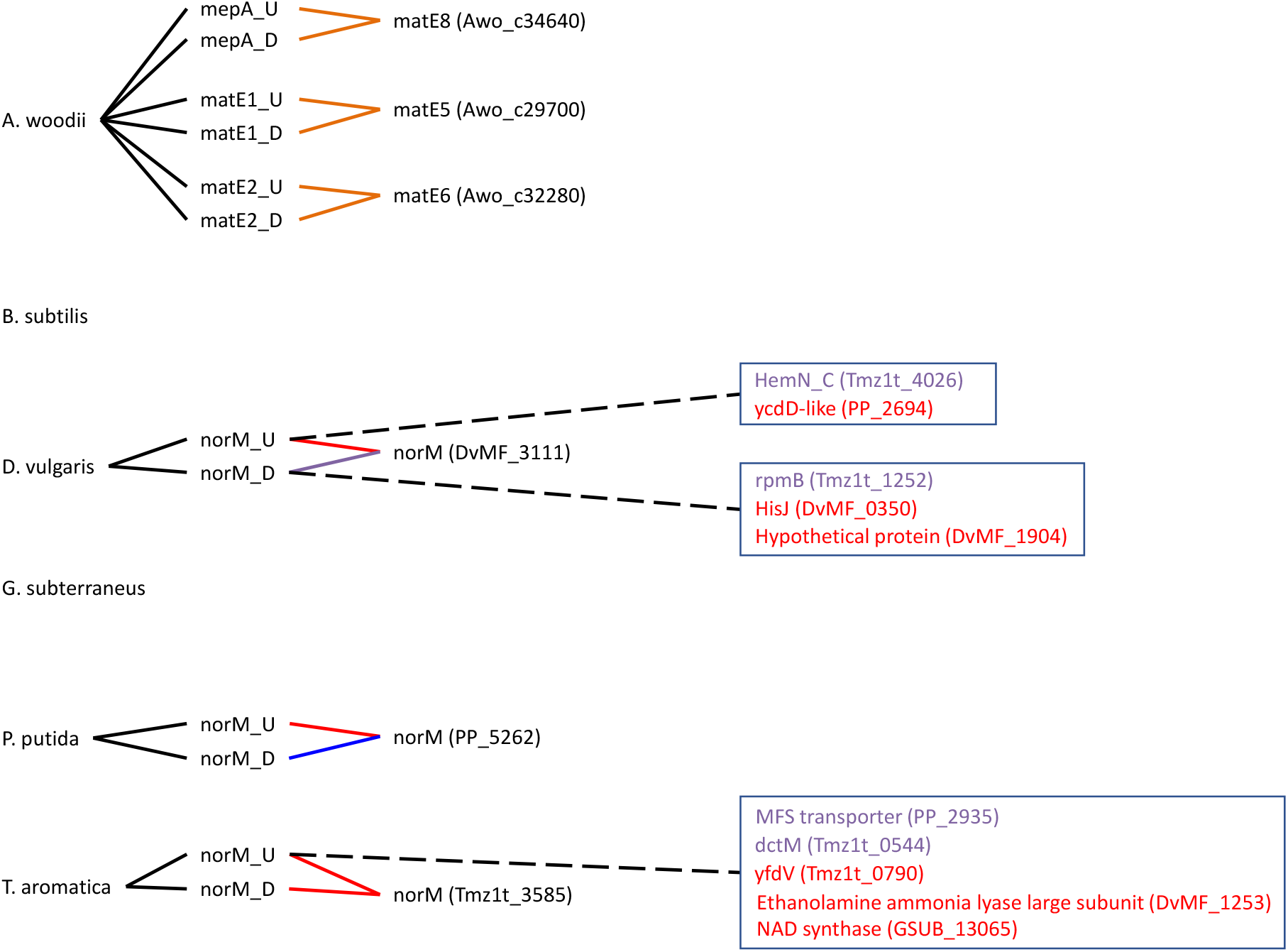
Visual representation of the primers designed in this study targeting the multidrug and toxic (compound) extrusion (MATE) genes and all targets they bind to. The type of line connecting the primer and the target indicate the homology of binding (solid line 100%, dashed line 90-99.9%, dotted line 80-89.9%). The colour coding is used to indicate the melting temperature range divided into 5 °C increments (light green ≤49.9 °C, orange 50-54.9 °C, blue 55.0-59.9 °C, purple 60.0-64.9 °C, red 65.0-69.9 °C and dark red ≥70.0 °C). A black line connecting to the gene targets grouped into a box indicates the homology of all genes in the box and the text colour represents the melting temperature.

**Figure 5.**
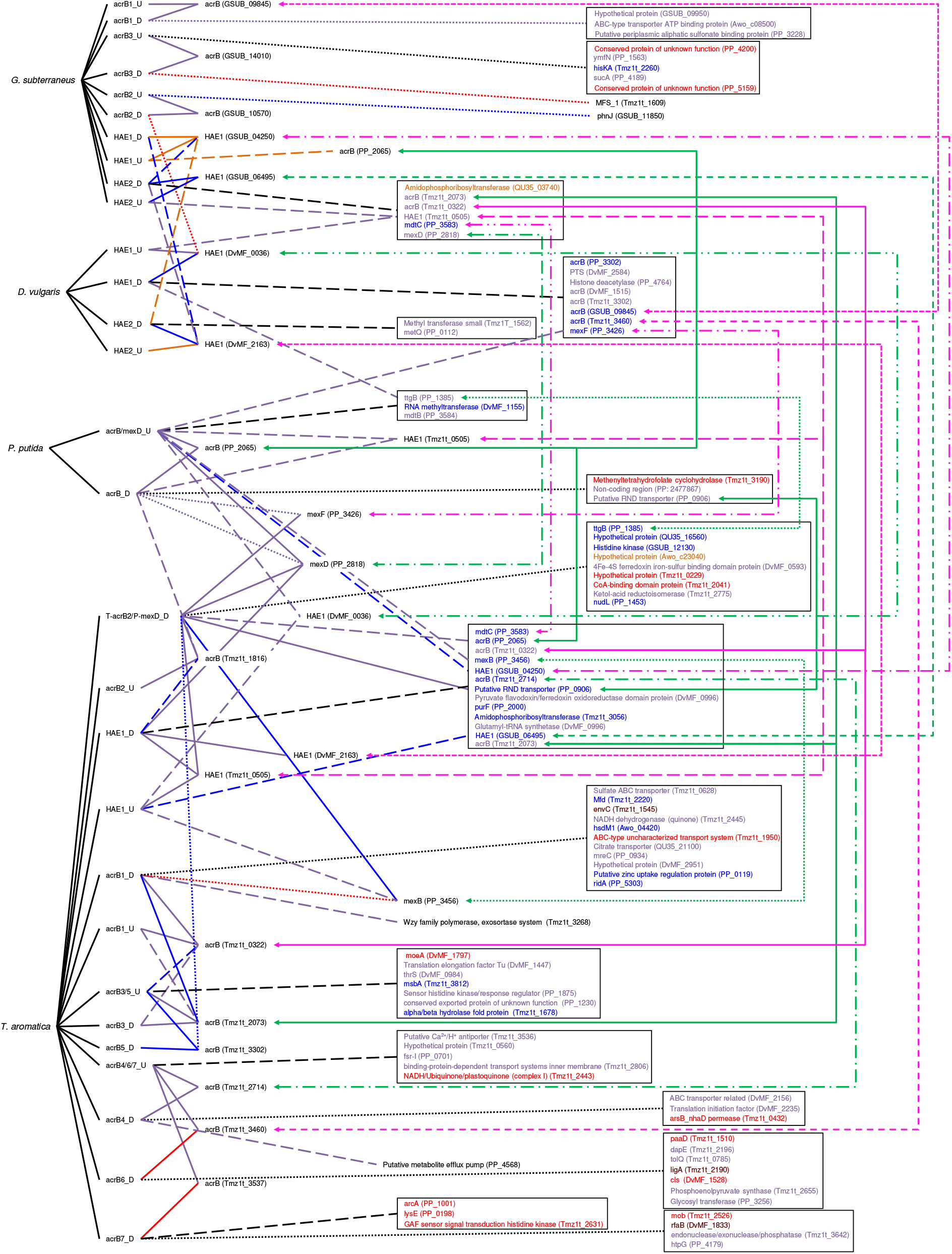
Visual representation of the primers designed in this study targeting the resistance-nodulation-cell division (RND) genes and all targets they bind to. The type of line connecting the primer and the target indicate the homology of binding (solid line 100%, dashed line 90-99.9%, dotted line 80-89.9%). The colour coding is used to indicate the melting temperature range divided into 5 °C increments (light green ≤49.9 °C, orange 50-54.9 °C, blue 55.0-59.9 °C, purple 60.0-64.9 °C, red 65.0-69.9 °C and dark red ≥70.0 °C). A black line connecting to the gene targets grouped into a box indicates the homology of all genes in the box and the text colour represents the melting temperature. Green and pink lines on the right link gene targets which have been duplicated to simplify visual representation.

This work illustrates the difficulty of making a universal primer set for accessory/character genes compared to core genes. The challenge is further highlighted working with genes that frequent mobile genetic elements thus providing opportunity for increased divergence. To illustrate the issues we observed and our findings, we discuss a few key examples highlighting certain trends and difficulties which were discovered as a result of this primer design work.

### Off target unintended primer binding

To begin, an example showing the potential of a primer designed for one target can identify other MDREPs in another species is discussed. The upstream *norM* primer targeting *T. aromatica* has a relatively high annealing temperature on its target sequence (65.6 °C) owing to its high GC content of 75%. This primer has three unintended binding targets, all with a binding homology of 90-99.9% (Fig. 4). Two of these sites are located within *T. aromatica* and target *dctM* (Tmz1t_0544) and *yfdV* (Tmz1t_0790). *dctM* is a tripartite ATP-independent periplasmic transporter which falls under the C4-dicarboxylate transport system classification according to KEGG while *yfdV* is an auxin efflux carrier but is only a hypothetical protein with general function predicted and thus could be a more general transporter. The third unintended target is a conserved membrane protein of unknown function in *P. putida* (PP_2935) but is provisionally in the MFS superfamily. The annealing temperatures of these locations are at or just below the intended annealing temperature. This illustrates a clean example of a primer targeting a sequence of nucleotides encoding what may be a conserved region of certain transporter genes. It is important to note that although these primers are detecting these unintended targets *in silico*, in each case only a single primer is binding and therefore no double stranded PCR product would be formed *in vitro.* In practice, these unintended single primer interactions will only affect PCR amplification by reducing the availability of the primers (i.e. primer efficacy) to find the intended sequence. Significant amounts of off-target binding interactions will reduce the availability of the primers to bind to the intended target sequence which decreases the amount of PCR amplicon production. This reduction in primer availability has additional implications for interpreting qPCR data as it may result in an underestimation of gene copies.

A more complicated example of unintended primer binding interactions is the downstream *qacA* primer for *D. vulgaris* (Fig. 1). In this example, the downstream *qacA* primer has an annealing temperature of 55.4 °C and three unintended binding sites, none of which are in the *D. vulgaris* genome. The primer binds at the same or lower annealing temperature to *pcrA* (QU35_03810), an ATP-dependent DNA helicase, and a succinyl-CoA synthetase alpha subunit (QU35_08905), both in *B. subtilis* and cupin gene (GSUB_03070) in *G. subterraneus.* This example illustrates undesired targeting which do not provide any sort of additional information such as potentially conserved sequences (domains) of efflux pumps or identifying unannotated/misannotated genes of similar function. It is unclear what the trait(s) of these genes is contributing towards being targeted by this primer.

Finally, we discuss the unintended targeting of three different primers, two of which were constructed with degenerate bases and intended to target multiple sequences in different species. The two degenerate primers are the upstream *acrB/mexD* targeting both genes in *P. putida* and the downstream primer targeting *acrB* in *T. aromatica* and *mexD* in *P. putida.* The single target primer is the downstream *acrB* for *P. putida* which complements the *P. putida acrB/mexD* primer. For the intended targets, all primers anneal with 100% homology. Due to the degenerate nature of two of these primers, there is a significant increase in the unintended binding targets compared to other primer sets (Fig. 5). Interestingly, the unintended binding locations of the upstream *acrB/mexD* primer are exclusively between 90-99.9% sequence homology while the *P. putida* downstream *acrB* primer has a mix of high homology and low homology binding sites. Some of the recurring unintended targets are the other *mex* genes, specifically *mexB* and *mexF*, both of which are detected by the *acrB/mexD* upstream and downstream primers with homologies of 90-99.9% (with the exception of *mexB*, which unintentionally has 100% homology with the *T. aromatica/P. putida mexD* downstream primer). The *T. aromatica*/*P. putida mexD* downstream primer also has 100% homology matches with *mexF* (PP_3426) and a putative RND transporter (PP_0906) (Fig. 5). Many of the unintended targets of this degenerate downstream primer are hypothetical proteins and all have annealing temperatures of 50-59.9 °C (a minimum of 5 °C below the melting temperature of the intended target). Of the unintended targets of the upstream *P. putida acrB/mexD* primer, two are hydrophobe/amphiphile efflux-1 (HAE1) genes (DvMF_0036 and Tmz1t_0505), one is an uncharacterized RND transporter in *G. subterraneus* (GSUB_04250) and the final target is the *ttgB* gene (PP_1385) in *P. putida*, which codes for a probable membrane efflux pump transporter protein.

This primer set illustrates the ability of the primers designed from an MSA of multiple genes (11 annotations in total) being able to target and locate other similar genes both within the intended target species and in other unrelated species owing partially to the degenerate bases present. Most hits are located within the two intended species with only two hits occurring outside, HAE1 in *D. vulgaris* and an RND transporter in *G. subterraneus.* It is important to note that RND primers are the only primers to have both upstream and downstream primers binding simultaneously to the same target, potentially resulting in unintended PCR amplicons. This occurs in four different genes: *mexF* (PP_3426), *ttgB* (PP_1385), *HAE1* (Tmz1t_0505) and *mexB* (PP_3456) (Fig 5). A fifth gene is targeted by multiple primers (putative RND transporter, PP_0906), however this gene is only targeted by downstream primers which bind to the identical location and thus cannot produce a PCR amplicon. Perhaps unsurprising, the primers targeting *mexD* in *P. putida* also bind to and would produce a PCR amplicon on *mexB* (PP_3456) and *mexF* (PP_3426), suggesting that these genes have a homologous domain which influenced the MSA alignment of the *mexD* genes. The other two unintended targets with multiple primer attachments target a *ttgB* (PP_1385) gene in *P. putida* which encodes for a probable efflux pump and an HAE1 gene in *T. aromatica* (Tmz1t_0505) which belongs to the acriflavine resistance protein B family (efflux pump) according to KEGG. It becomes clear from this example that the degenerate bases allow for an increase in unintended target locations, and unlike the other examples, these primers will produce actual PCR amplicons, disrupting any potential qPCR or downstream analyses.

An issue resulting from our Primer Design A approach (i.e. designing with degenerate bases), we developed an alternative method (Primer Design B) to create primer sets which preferentially target different locations (i.e. potentially unconserved locations), but prioritize maintaining identical amplicon sizes across all intended targets. As illustrated with the RND primers, Design A can lead to an increase in unintended binding locations due to the exponential increase in primer sequences. Each degenerate base increases the number of unique sequences present in the primer mix and may create a primer which has no intended target sequence meaning the chance of unintended binding increases. To illustrate, take the following example for the upstream primer targeting the *acrB* and *mexD* genes in *P. putida*:

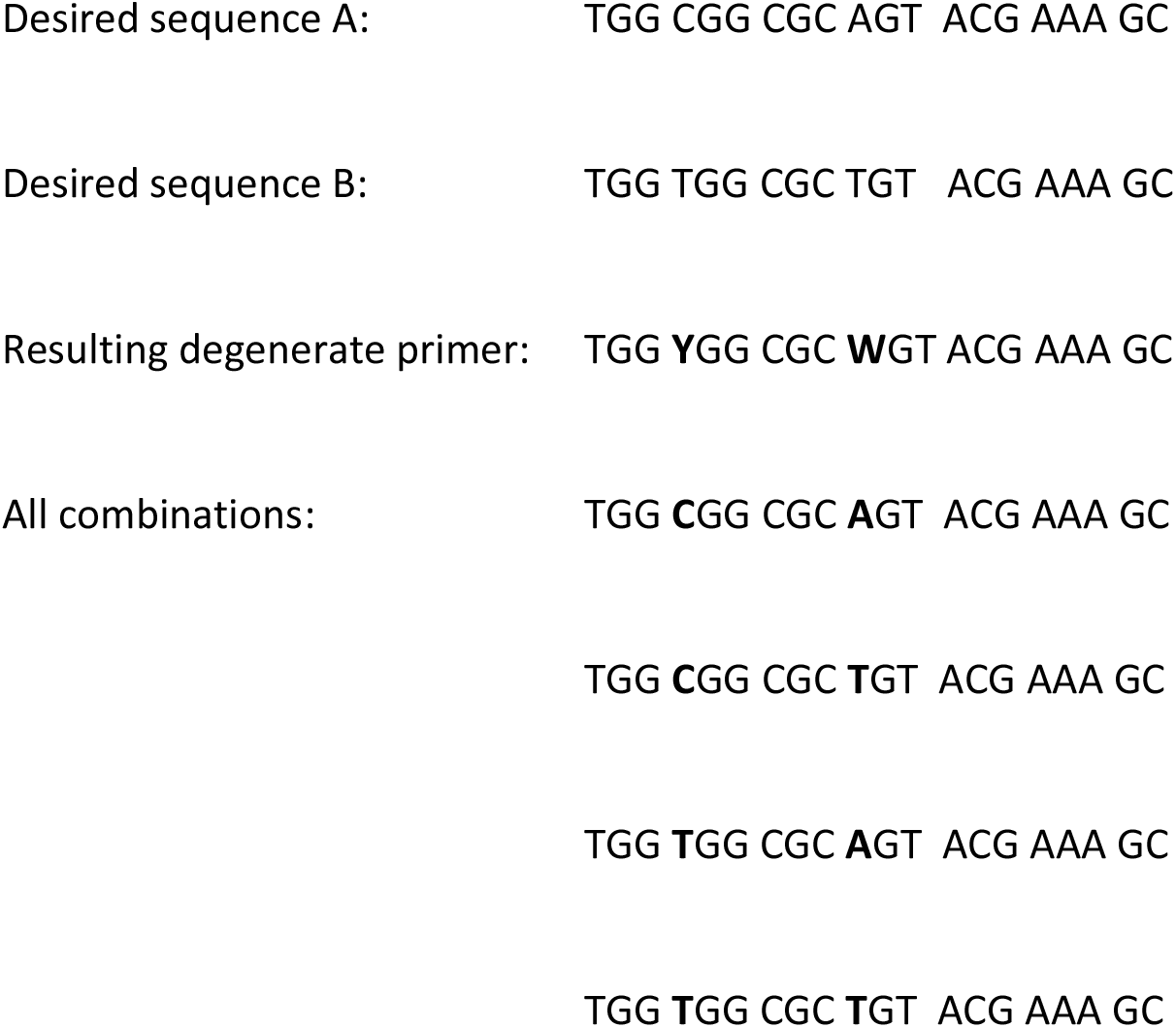

From this simple example using two degenerate bases coding for only two nucleotides, the resulting degenerate primer consists of a mixture of four primers, two of which do not code for any intended sequence. Rather, a more precise approach is to treat each gene as its own template, design primers to create the same amplicon size and have similar melting temperatures and subsequently combine all the upstream primers into a single mixture in equal proportions. In this way, the number of primers is kept to a minimum and every primer has a desired target.

The unintended specificity of these primers is of note when considering that all the primers are 18-23 nucleotides in length and therefore target a sequence of six to seven amino acids. It must be understood that not all MDREP genes would be detected using this approach but it could provide a means of improving our understanding of conserved regions, and as this work illustrates, the conserved amino acid region may be as short as six to seven amino acids and still provide relatively high accuracy for gene identification. It is clear from this that even a six amino acid sequence can have evolutionary pressure to convey similarity in overall protein structure and/or function.

Using the approach B (i.e. mixing each unique primer together without degenerate bases), it allows the “universal” primer mix to always be in flux and be improved as new primers can always be added, until such point the number of primers present negatively affects a specific primer’s ability to bind the correct sequence. A consideration with Approach B is that because the primers do not always target conserved regions, the primers may have unintended targets elsewhere in the genome(s), entirely unrelated to the target sequence. As a result (and as is for good practice), primer sequences developed using either approach should be tested against the target genome(s) and explore all unintended binding locations should be identified and accounted for during PCR protocol development.

### MDREP primers as compared to universal 16S rRNA primers

In contrast to the successful universal primer designs targeting the 16S rRNA gene, the issues discussed here highlight the difference between targeting core genes (essential for replication), character genes (those defining the type of metabolism) and accessory genes (genes which may provide improved fitness under specific conditions). As we improve techniques and apply genetic screening to more and more health (e.g. infection, disease, etc.) and economically (e.g. agriculture, bioremediation, etc.) issues, the more relevant genetic targets we will discover. Logically, the more specific the target, the more meaningful the presence/absence and quantity present becomes, but the further away from the core genes moving towards characteristic and accessory gene we must move. Accessory genes (and to a lesser extent characteristic genes) have less evolutionary pressure on them which allows for higher rates of mutation and variability on the nucleotide level. Additionally, as many accessory genes are or can be located on mobile genetic elements, they become subject to the nucleotide biases and codon usages present in their current host. Furthermore, the mechanism of the movement may affect the gene’s availability or expression levels. The fitness of an accessory gene is dependant on environmental pressures and are subject to pulses of challenge such as short periods of exposure to biocide challenges as is typical for pipeline antimicrobial treatments or antibiotic courses.

For comparisons, the percent identities of the target genes were calculated using the multiple sequence alignment (MSA) for each respective gene. The scores were calculated using the equation:

> Percent Identity = (matches * 100)/length of MSA (including gaps)

Each MSA was calculated using the conditions mentioned in Methods. Only a single representative gene for each super family was selected as an indication of the variability within each superfamily. The average scores of the sequence identities are listed in Table 3 and a complete list for each genes scores are provided in supplementary data (Supplementary Tables 2-6). The ABC superfamily has been omitted due to their low gene counts across the six representative species.

**Table 3.**
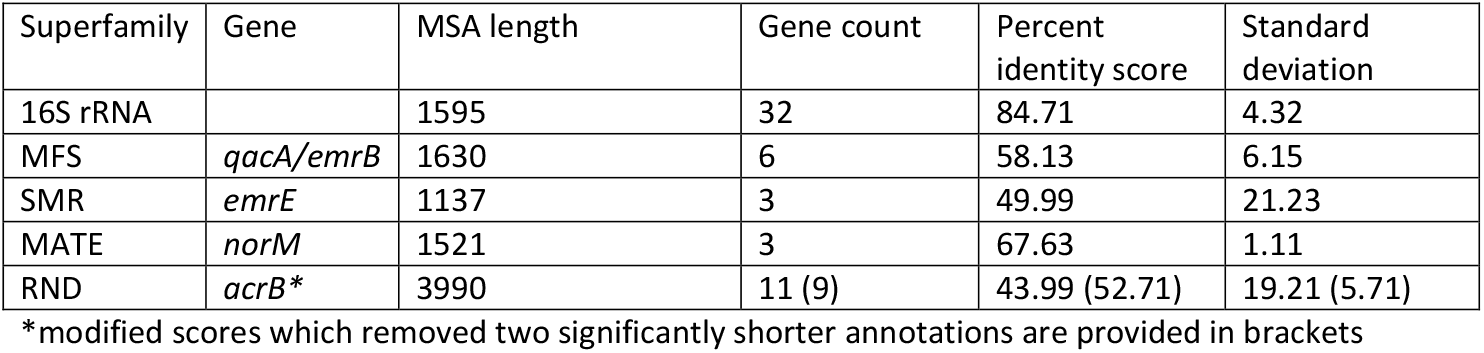
Average scores of percent identities for genes chosen to represent four superfamilies

From the percent identity scores, it is obvious the 16S rRNA gene is more conserved compared to any of the efflux pumps. The highest score belongs to the *norM* gene which, among the annotated copies had very high homology (67.63%) while *acrB* and *emrE* had low scores (43.99 and 49.99% respectively). The *acrB* score is skewed from two annotated copies being significantly shorter than the MSA, producing percent identity scores of 5.26% and 4.24%. Removing these two copies produces a score of 52.71% ± 5.71, which brings the score for *acrB* to be closer in line with *qacA/emrB* and *emrE* scores.

These scores reflect the relative simplicity of designing primers for highly conserved, core genes while nonessential, accessory genes are significantly more difficult due their more plastic nature. While they are less homologous, the identity scores suggest it is possible to design universal primers for these peripheral genes. However, these genes do require a different approach for primer design. Using approach A, which employs multiple sequence alignments, more conserved regions can be identified to target but using approach B, which employs unique primer sequences (avoiding the use of degenerate bases) will result in a primer mixture with higher accuracy and specificity. Both approaches will still be able to identify additional, related genes during *in silico* analyses.

The plasticity of nucleotide sequences of the accessory genes results from the improved fitness benefits only occurring under specific conditions which are not perpetually present. This allows for mutations to occur which would otherwise be impossible in core or character genes. Due to the non-conserved nature of MDREP genes and their mobility through and across genomes, the abundance and variance on the genetic level is erratic and unpredictable. Presence of same gene in two different species doesn’t allow for the determination of the direction of flow or origin of the gene. This *in silico* approach has the potential to be exploited as a means to investigate evolutionary branching in these genes and in the case of MDREP genes, potentially shedding light on how these genes are selected for and allow for the identification of other clinically relevant efflux pumps yet to be discovered.

### Lessons learned

An overlying issue with primer design attempts such as this one is the often-poor annotation quality of our genomic data libraries, particularly for environmental (or more generally any non-clinical) species. The abundance of putative, predicted or hypothetical proteins severely limits the ability to accurately find related genes to design primers for specific genes. To alleviate this issue, primers should be designed using a well annotated genome which is likely to be present in the environment where the primers will ultimately be used. In this way, the primers can be designed with higher confidence and as shown here, can be used to probe genomes with lower annotation quality and aid in identifying the desired targets there.

As with all scientific endeavours, it behooves the scientist to keep in mind the ultimate goal of the primers. Here, we attempted to design primers for the same genes across six different genomes to eventually combine them into a “universal” primer mix for the desired target. In these situations, unintended binding becomes a larger issue as the amount of a specific primer is already reduced with respect to the final primer concentration and any off-site binding could result in false negatives during PCR amplification. If the objective is to probe a mixed sample for the presence of a specific gene *(e.g.* for clinical or environmental screening), the use of degenerate primers becomes risky as it may lead to false positives or negatives (should the mixture of primers be too complex and the competitive binding of the primers prevents correct primer binding). Alternatively, in exploratory science such as the attempt explained here, the degenerate primers can increase the ability of the primers to detect additional genes not identified in the annotations. There is the potential that of many of the detected genes coding for hypothetical proteins are truly efflux pumps of some nature and through this approach we can add additional evidence towards these predicted proteins. This would require a more targeted investigation of the genes identified in this manner such as comparing the amino acid sequences of the predicted genes to known proteins and determining whether they truly would produce efflux pumps.

Unlike the 16S rRNA gene family, designing universal primers for mobile, accessory genes is particularly difficult. Of note from this approach is the utility and the potential of *in silico* analysis of primers designed for less conserved genes and their potential to aid or facilitate improved annotation in lesser studied organisms.

The choice between which of the two primer design approaches should be employed becomes a decision based on the homology or conservation of the nucleotide sequences of the target genes. To facilitate the decision, percent identity scores should be calculated for all annotated copies of the desired target gene. From the 16S rRNA and MDREP gene examples shown here, we suggest a cut-off value of 75%, where identity scores above 75% should use primer Design A (employing MSA and degenerate bases), while identity scores below 75% would be more effective if primers were designed using Design B (unique sequences with variable binding locations). This should reduce the amount of unintended binding locations and overall improve the efficacy of primer binding in PCR and qPCR applications.

## Conclusion

This work illustrates the benefits and shortcomings of two different primer design approaches. First, the use of multiple sequence alignments (MSAs) to locate conserved regions of the nucleic acid lends itself towards creating primers with degenerate bases and ensures uniform PCR amplicon size. With the degenerate bases, these primers are more likely to have unintended binding and lower primer efficiency during thermocycling. However, these primers are more likely to reveal the presence of the desired gene(s) in poorly annotated genomes when used in a *in silico* method.

The second primer design approach creates primers from individual genes without targeting conserved regions of the MSA while controlling for melting temperature and amplicon size to ensure all primers designed in this way are compatible. This approach allows for more stringent thermocycling conditions and reduces the amount of unintended primer binding locations (thus improving detection rates *in vitro)*, but this approach is less likely to reveal similar or the same genes in mixed environments.

Overall this work highlights how extremely important it is to appreciate the false discovery rates resulting from the chosen primer design approach and the subsequent ramifications in interpretations when it comes to defining the answer to one’s goal.

## Acknowledgements

This work was supported by Genome Canada through a Large Scale Applied Research Project grant. D.C.B. was supported by a PhD scholarship from NSERC.

